# Endothelial-pericyte interactions activate insulin signaling and its implications for blood-brain barrier dysfunction in Alzheimer’s disease

**DOI:** 10.1101/2025.11.24.690270

**Authors:** Zengtao Wang, Vaishnavi Veerareddy, Paulina M. Eberts, Samira M. Azarin, Karunya K. Kandimalla

**Affiliations:** Department of Pharmaceutics and Brain Barriers Research Center, University of Minnesota, College of Pharmacy, Minneapolis, MN; Department of Chemical Engineering and Materials Science, College of Science and Engineering, University of Minnesota, Minneapolis, MN

**Author notes:** Corresponding author: Karunya K. Kandimalla, Department of Pharmaceutics and Brain Barriers Research Center, College of Pharmacy, University of Minnesota, Minneapolis, MN, 55455, USA. Global PK/PD/Pharmacometrics, Eli Lilly and Company, Lilly Corporate Center, Indianapolis, Indiana, 46285, USA.

**Keywords:** TIMP1, MMP, insulin signaling, exosomes

## Abstract

**Purpose:** This study aimed to investigate how pericyte degeneration contributes to BBB disruption in Alzheimer’s disease, focusing on the roles of insulin signaling and the imbalance between matrix metalloproteinases (MMPs) and endogenous tissue inhibitors of MMPs (TIMPs).

**Methods:** We employed an in vitro BBB model by co-culturing brain-specific microvascular endothelial-like cells (iBMECs) differentiated from human induced pluripotent stem cells (hiPSCs) and primary human brain vasculature pericytes (hBVPs). Protein expression under solo- versus co-culture conditions was assessed by western blot. MMP enzymatic activity in the culture media was measured by fluorometric assay. Exosomes were isolated from conditioned media and brain derived neurotrophic growth factor (BDNF) concentrations were determined using ELISA assays.

**Results:** TIMP1 and collagen-IV expression was significantly increased in co-cultured BBB endothelial cells and pericytes compared to solo-cultures. However, a greater effect was observed in cells co-cultured for 2 days than 7 days. Elevated TIMP1 in co-culture media significantly inhibited MMP activity. The AKT and ERK pathways were activated in both cell types after 7 days of co-culture, and the ERK signaling mediated TIMP1 upregulation in endothelial cells. BDNF was significantly enriched in exosomes isolated from co-culture media on the abluminal side compared to the solo-cultures. Endothelial cells also protected pericytes from accumulation of toxic amyloid-beta 42 by downregulating low density lipoprotein receptor-related protein 1 (LRP1) expression.

**Conclusions:** These findings provide mechanistic insights into BBB disruption due to pericyte degeneration and highlight the important role of BBB insulin resistance in causing cerebrovascular dysfunction in AD.

## 1. Introduction

Cerebrovascular dysfunction is prevalent in 80% of Alzheimer’s disease (AD) patients without vascular dementia[1] and is closely linked to cognitive decline in AD. Impairment of the blood-brain barrier (BBB), a cellular monolayer lining of the cerebrovascular lumen, is regarded as a significant contributor to cerebrovascular dysfunction in AD. The BBB is primarily composed of brain endothelial monolayers with strictly regulated paracellular and transcellular passage of molecules, achieved by tight junctions and restrictive intracellular vesicular trafficking, respectively. Additionally, polarized distribution of receptors/transporters on the luminal versus abluminal side of the BBB maintains the directionality and specificity of BBB trafficking and signaling.

However, BBB functions are not handled solely by the endothelial cells; instead, interactions with cellular constituents of the neurovascular unit (NVU) are critical to maintaining BBB functions. The NVU is formed by the specialized extracellular matrix (ECM) of the basement membrane connecting endothelial cells with neighboring neuroglial cells like pericytes and astrocytes. Key molecular components of the ECM secreted by both endothelial and neuroglia cells include collagen, laminin and fibronectin. They provide the structural support and mediate the cell-cell interactions that ensure the functionality of a mature BBB. Notably, disruption of ECM has garnered considerable attention in AD research, primarily because it contributes to various pathologies, such as inhibiting synaptic transmission, promoting amyloid beta (Aβ) accumulation and inducing inflammation[2].

Enzymes of the matrix metalloproteinases (MMP) family are proteinases that regulate the ECM remodeling. MMP has been shown to disrupt the BBB by degrading both tight junctions and ECM proteins in various brain disorders including neuroinflammation[3], stroke[4], and multiple sclerosis[5]. In AD, it was found that Aβ upregulated MMP expression and activity, which caused tight junction damage and BBB leakage[6,7]. MMP enzyme activity is modulated by tissue inhibitors of metalloproteinases (TIMPs), which are endogenous inhibitors of MMP. The TIMPs comprise four members known as TIMP1-4. The balance between MMP and TIMP proteins are critical in preserving BBB integrity in various diseases. For instance, TIMP1 knockout mice exhibited exacerbated BBB disruption and neuronal apoptosis during focal cerebral ischemia compared to wild-type mice[8]. Conversely, in a mouse model of intracerebral hemorrhage, TIMP1 treatment was shown to alleviate BBB disruption and improve neurological performance[9]. Nevertheless, the protective function of TIMP in preventing BBB disruption in AD has not been characterized.

Pericyte degeneration is a pivotal characteristic and a major contributor to cerebrovascular dysfunction in AD[10–12]. It is well established that pericytes regulate blood flow by controlling microvascular dilation. In the AD brain, it was observed that Aβ constricted the capillaries specifically at pericyte locations, leading to a reduction of cerebral blood flow[13]. Pericyte degeneration has also been linked to downregulation of endothelial tight junction expression, including ZO-1 and occludin, resulting in BBB disruption in both AD animals and patients[14,15]. These findings underscore the critical role of pericyte degeneration in cerebrovascular dysfunction in AD. However, the impact of pericyte loss on the ECM of the BBB, has not been reported yet.

Our previous transcriptomic analysis indicated that insulin stimulation upregulated the gene expression of all TIMPs[16]. Further, we recently showed that pericytes influence changes in the transcriptome of BBB endothelial cells, thereby providing protection against insulin resistance[17]. Thus, we hypothesized that the interaction between endothelial cells and pericytes might contribute to BBB integrity by regulating endothelial TIMP expression through the insulin signaling pathway. In this study, we tested this hypothesis using an in vitro BBB model, constructed by co-culturing human induced pluripotent stem cells derived brain-specific microvascular endothelial-like cells (iBMECs) with primary human brain vasculature pericytes (hBVPs). Further, we explored the protective effects of BBB endothelial cells on pericytes and their implications in AD.

## 2. Materials and Methods

### 2.1. Cell culture

Human induced pluripotent stem cells (hiPSCs; IMR90-4 line, WiCell) were differentiated into iBMECs following established protocols as described in previous publications[17–19]. On day 8 of differentiation, cells were seeded onto plates coated with a mixture of collagen IV and fibronectin (CN IV-FN) to selectively enrich endothelial cells.. After 1 to 2 hours, unattached cells were removed, and the remaining cells were rinsed once with Dulbecco’s Phosphate-Buffered Saline. The enriched endothelial population was then transferred onto 24 mm CellTreat® inserts (CELLTREAT Scientific Products, Pepperell, MA) at a seeding density of 1 × 10^6^ cells/cm^2^. Cultures were maintained for one day in endothelial cell medium to promote barrier formation. The endothelial cell medium consists of human endothelial serum free medium (Thermo Fisher Scientific, Waltham, MA) supplemented with 1% fetal bovine serum (Thermo Fisher Scientific), 20 ng/ml basic fibroblast growth factor (bFGF) (Thermo Fisher Scientific), and 10 μM retinoic acid (Millipore Sigma, Burlington, MA). After approximately 24 hours, the medium was replaced with endothelial cell medium lacking retinoic acid and bFGF. Frozen vials of human brain vascular pericytes (hBVPs; ScienCell, Carlsbad, CA) (passage 1; >5 × 10⁵ cells) were quickly thawed and plated into gelatin-coated T-75 flasks (Thermo Fisher Scientific) containing pericyte growth medium (ScienCell, Carlsbad, CA). Once hBVPs reached 90-95% confluence, they were detached using trypsin and then seeded at a concentration of 5,000 cells/cm^2^ onto the bottom of gelatin-coated 6-well plates. Two experimental designs were employed in the current study. In the 2-day co-culture model, purified iBMECs were initially cultured alone on cell culture inserts for 7 days to establish a mature barrier. Subsequently, pericytes were introduced and co-cultured for 2 days. In the 7-day co-culture model, iBMECs were co-cultured with pericytes together for the entire duration of 7 days to form a mature barrier. Co-cultures were established by placing CellTreat® inserts carrying iBMECs into 6-well plates pre-seeded with pericytes. The culture medium was then replaced with endothelial cell medium to initiate a non-contact in vitro BBB model, which was maintained at 37°C in a humidified incubator with 5% CO₂. The hCMEC/D3 cell line was kindly provided by P-O Couraud (Institut Cochin, France). Cells were grown under 5% CO_2_ at 37°C as reported previously[20]. The endothelial cell medium consists of basal medium-2 (Lonza, NJ) containing 5% of fetal bovine serum. Various supplements including 1 ng/ml human basic fibroblast growth factor (PeproTech, NJ), 1% chemically defined lipid concentrate (Gibco, NY), 10 mM HEPES, 5 µg/ml ascorbic acid, 1.4 µM hydrocortisone, and 1% penicillin-streptomycin (MP Biomaterials, OH) were added to the cell culture medium.

### 2.2. Western blot

After co-culture, both iBMECs and hBVPs were washed three times with PBS and lysed with RIPA buffer containing protease and phosphatase inhibitors (Sigma-Aldrich, St. Louis. MO). Total protein concentrations in the lysates were determined by bicinchoninic acid (BCA) assay (Pierce, Waltham, MA). The lysates (25 µg protein per lane) were loaded onto 4-12% Criterion XT precast gels and proteins were separated by SDS-PAGE under reducing conditions (Bio-Rad Laboratories, Hercules, CA). The proteins were then electroblotted onto a 0.45 µm nitrocellulose membrane, blocked with 5% nonfat dry milk protein (Bio-Rad Laboratories, Hercules, CA), and incubated overnight at 4 °C with primary antibodies against TIMP1 (1:500, Cell Signaling Technology, Danvers, MA), Vinculin (1:2000, Cell Signaling Technology, Danvers, MA), p-AKT (1:1000, Cell Signaling Technology, Danvers, MA), p-ERK (1:1000, Cell Signaling Technology, Danvers, MA), total-AKT (1:1000, Cell Signaling Technology, Danvers, MA), total-ERK (1:1000, Cell Signaling Technology, Danvers, MA), GAPDH, claudin-5 (1:1000, Cell Signaling Technology, Danvers, MA) and collagen-IV (1:1000, Thermo Fisher Scientific). Afterwards, the membrane was incubated with IR-dye conjugated secondary antibody (1:2000, LI-COR Inc, Lincoln, NE) for 1 h at room temperature. All blots were derived from the same experiment and were processed in parallel. Immuno-reactive bands were then imaged (Odyssey CLx; LI-COR Inc, Lincoln, NE) and the band intensities were quantified by densitometry (Image StudioTM Lite Software, LI-COR Inc, Lincoln, NE).

### 2.3. MMP activity assay

The general activity of MMP enzyme was quantified using the MMP Activity Assay Kit (Fluorometric-Green) (ab112146, Abcam, Cambridge, UK) following the instructions provided by the manufacturer. Briefly, cell culture media were collected from both luminal and abluminal sides of the iBMEC monolayer and then incubated with the fluorescence-resonance-energy-transfer–based MMP green substrate for 1 hour. The fluorescence intensity was determined on a fluorescent plate reader (Molecular Devices, San Jose, CA) at E_x_/E_m_ = 490/525 nm. In the inhibitor study, the iBMECs were treated with 10 µM Trametinib for 6 hours. hCMEC/D3 cells were pretreated with Trametinib (10 µM) for 24 hours followed by insulin (100 nM) stimulation for 30 minutes. The results were presented as fold change compared to solo-culture or cells untreated with inhibitors.

### 2.4. Flow cytometry

The FITC-labeled Aβ_42_ peptides (F-Aβ_42_) were purchased from AAPPtec, LLC (Louisville, KY) and were reconstituted as described in our previous publication[21]. After 7-day co-culture, iBMECs were removed from hBVPs and media were aspirated. The solo- and co-cultured hBVPs were then incubated with 1 µM F-Aβ_42_ for 1 hour. For inhibitor studies, the hBVPs were pretreated with caveolae-mediated endocytosis inhibitors Nystatin (2.5 µM, Fisher Scientific) or Filipin-III (1 µg/ml, Cayman Chemical) for 30 minutes before F-Aβ_42_ addition. After the treatment, the cells were washed twice with ice cold PBS and then subjected to trypsinization. The dislodged cells were using ice cold PBS, fixed with 4 % PFA solution and analyzed for intracellular fluorescence using BD FACSCalibur™ flow cytometer. The F-Aβ_42_ intracellular fluorescence intensities were measured using a 488 nm laser fitted with 530/30 filter. Data was acquired with BD CellQuest™ Pro and analyzed using FlowJo software. The F-Aβ_42_ uptake was represented as histograms of intracellular fluorescence intensities. All sample analyses were performed within one hour from the completion of the experiment.

### 2.5. Exosome isolation and characterization

To isolate exosomes from the culture media, iBMECs were cultured in regular medium containing Gibco^TM^ exosome-depleted FBS (Fisher Scientific). Total exosome isolation reagent (from cell culture media) was obtained from Thermo Fisher Scientific.

1.2mL of culture media was mixed with 0.6mL of separation reagent for each sample, and the mixture was incubated overnight at 4 °C. After incubation, the samples were centrifuged at 4 °C for 1 h at 10,000 × g. Finally, the precipitated exosomes were dispersed in 25µL of 1X PBS. The exosome suspension was diluted 1:200 for nanoparticle tracking analysis (NTA) using Nanosight LM-10 (Malvern Instruments Ltd., Malvern, UK), configured with a temperature controller and equipped with a 405 nm laser (near UV) inside the flow cell. The sCMOS camera recorded the Brownian motion of individual particles at a rate of 30 frames per second. Video of the moving particles was then obtained and processed using NanoSight’s nanoparticle tracking 2.3 image analysis software (NanoSight, USA) to evaluate their displacement in the x-y plane. This displacement data was subsequently utilized to calculate the diffusion coefficient, which in turn was used to determine the hydrodynamic diameter using the Stokes-Einstein equation. To lyse the exosomes, 10x RIPA buffer (ab156034, Abcam, Cambridge, UK) supplemented with a proteinase inhibitor (Sigma-Aldrich, St. Louis. MO) was added to the suspension and sonicated for 15 seconds. Western blot analysis was conducted using anti-CD9 antibody (1:1000, Cell Signaling Technology) to characterize the exosomes.

### 2.6. BDNF quantitative ELISA

The BDNF protein concentration in exosome lysates was determined using Human Free BDNF Quantitative ELISA Kit following the protocol provided by the manufacturer (R&D Systems, Minneapolis, MN, USA). Data were represented as pg/mL protein.

## 3. Results

### 3.1. TIMP1 expression increased in endothelial cell and pericyte co-cultures

Western blot analysis was conducted to examine TIMP1 expression in iBMEC and pericyte solo-cultures versus co-cultures. To evaluate the dynamics of endothelial-pericyte interactions, we implemented two experimental designs, with co-cultures maintained for either 2 days or 7 days (**Figure 1A**). As shown in **Figure 1B-E**, TIMP1 expression in iBMECs significantly increased when co-cultured with pericytes for both 2 days and 7 days. Interestingly, the fold-change was higher in the 2-day co-culture (6-fold increase) than the 7-day co-culture (2.5-fold increase). Pericytes also showed a significant 6-fold increase in TIMP1 expression after 2-day co-culture compared to the solo-culture (**Figure 1G, I**). However, there was no significant difference in pericyte TIMP1 expression between solo-culture and 7-day co-culture (**Figure 1H, J**). Given that TIMP1 is an endogenous inhibitor of MMP enzyme, we also determined the MMP activity in the culture media. It was found that the MMP activity was significantly decreased in the iBMEC culture media of 7-day co-cultures, specifically on the luminal side of the cell monolayer (**Figure 1F**).

**Figure 1.**
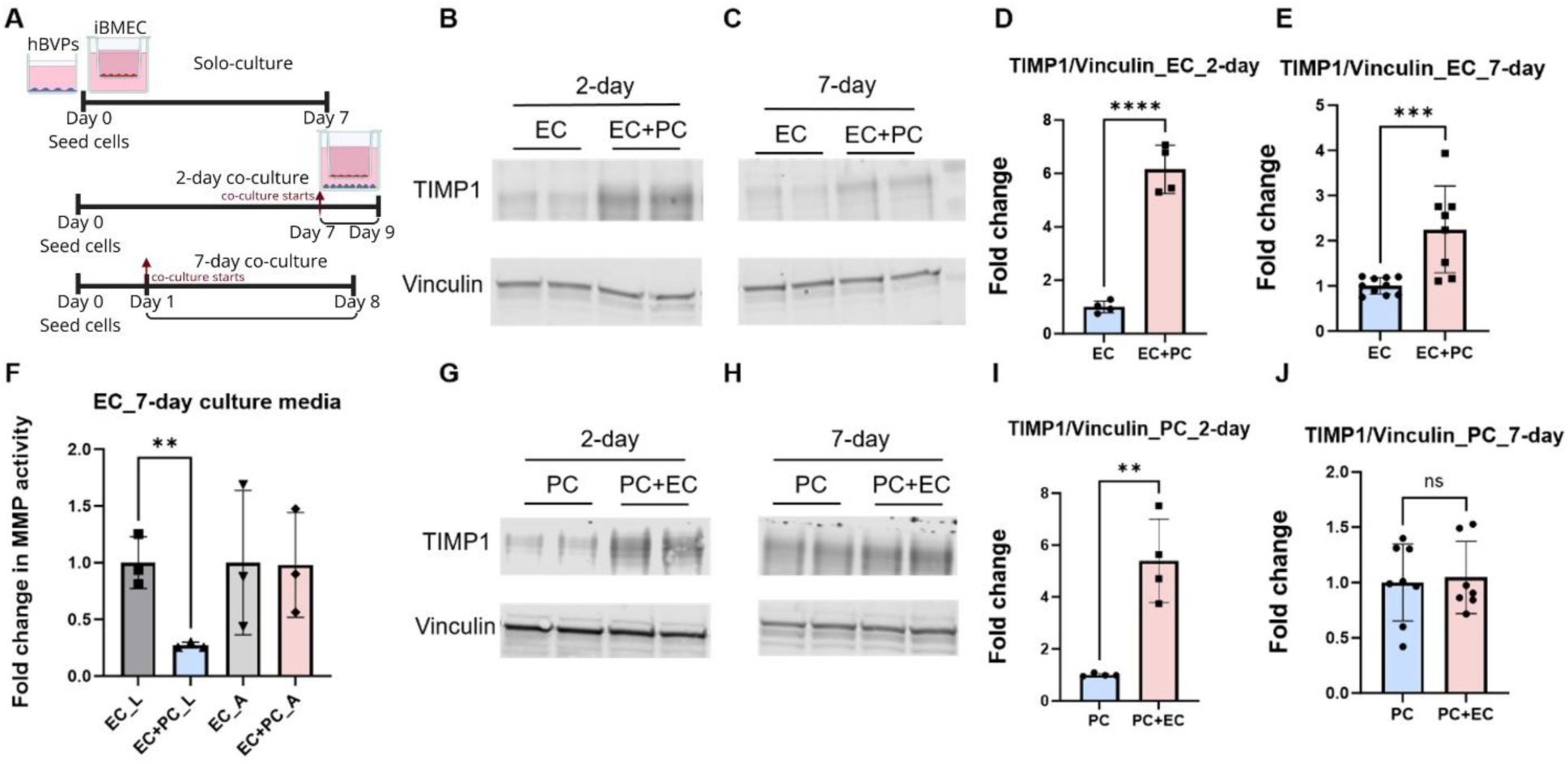
Increased expression of TIMP1 in co-cultured BBB endothelial cells and pericytes. **(A)** Experimental design. In 2-day co-cultures, iBMEC cells were cultured alone on cell culture inserts for 7 days to form a mature barrier, followed by co-culturing with pericytes for an additional 2 days. However, in the 7-day co-cultures, iBMEC cells were co-cultured with pericytes for 7 days to form a mature barrier. **(B-C)** Representative immunoblots showing TIMP1 and Vinculin expression in iBMEC cells with or without pericyte co-culture for **(B)** 2 days and **(C)** 7days. EC: endothelial cells, PC: pericytes. **(D & E)** Densitometric analysis of TIMP1 immunoblots normalized to Vinculin in (B) and (C). **(F)** MMP enzyme activity in culture media sampled from luminal (L) and abluminal (A) sides of EC solo-cultures and EC+PC co-cultures. **(G & H)** Representative immunoblots showing TIMP1 and Vinculin expression in pericytes with or without co-culture with iBMECs for **(G)** 2 days and **(H)** 7 days. **(I & J)** Densitometric analysis of TIMP1 immunoblots normalized to Vinculin in (G) and (H). Data in all the bar charts were presented as fold change compared to solo-cultures. (Mean ± SD, N represents the number of wells from three independent differentiation batches.). **p<0.01, ***p<0.001, ****p<0.0001, Student’s t-test.

### 3.2. Collagen-IV expression increased in endothelial cell and pericyte co-cultures

Collagen-IV is a major component of basement membrane of the BBB and is susceptible to degradation by activated MMP enzymes. Given the upregulation of TIMP1 expression and the decrease in MMP activity observed in the co-cultures, we next investigated the impact of co-culture on collagen-IV level in both iBMECs and pericytes. Our results demonstrated a significant increase in collagen-IV expression in iBMECs when co-cultured with pericytes for 2 days **(Figure 2A, C)** and 7 days **(Figure 2B, D)**. A similar trend was observed in pericytes. Specifically, in the 2-day co-cultures, collagen-IV expression in pericytes increased by 7-fold compared to solo-cultures **(Figure 2E, G)**. Co-culture with iBMECs for 7 days also significantly upregulated collagen-IV expression in pericytes, but the fold change was smaller than that in the 2-day co-cultures **(Figure 2F, H)**.

**Figure 2.**
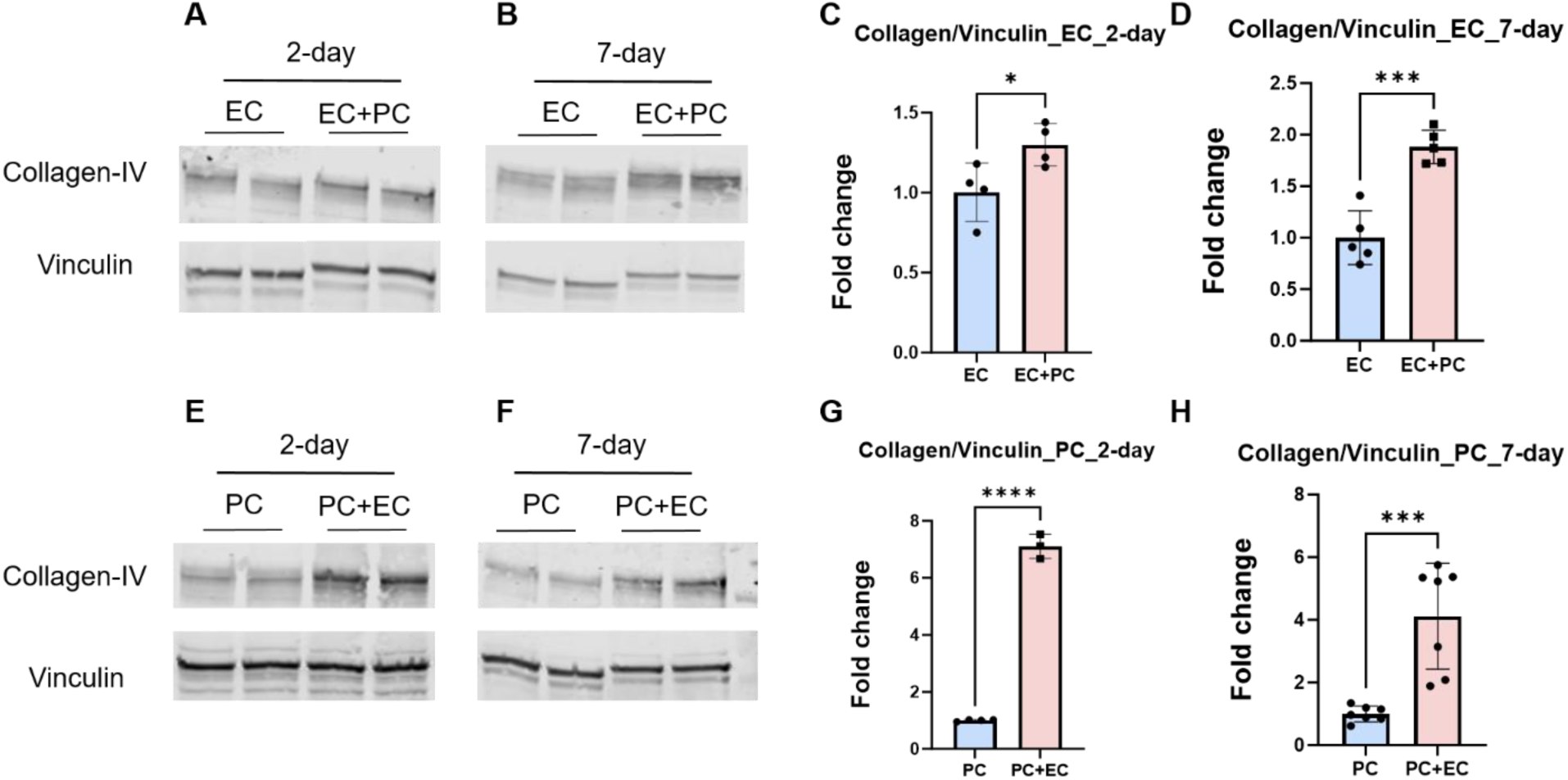
Increased expression of collagen-IV in co-cultured BBB endothelial cells and pericytes. **(A-B)** Representative immunoblots showing collagen-IV and Vinculin expression in iBMECs with or without co-culturing with pericytes for **(A)** 2 days and **(B)** 7 days. **(C-D)** Densitometric analysis of collagen-IV immunoblots normalized to Vinculin in (A) and (B). **(E-F)** Representative immunoblots showing collagen-IV and Vinculin expression in pericytes with or without co-culture with iBMECs for **(E)** 2 days and **(F)** 7 days. **(G-H)** Densitometric analysis of collagen-IV immunoblots normalized to Vinculin in (E) and (F). Data in all the bar charts were presented as fold change compared to solo-cultures. (Mean ± SD, N represents the number of wells from three independent differentiation batches). *p<0.05, ***p<0.001, ****p<0.0001, Student’s t-test.

### 3.3. AKT and ERK pathways were activated in endothelial cell and pericyte co-cultures

After characterizing the impact of co-cultures on TIMP1 and collagen-IV expression in BBB endothelial cells, we aimed to investigate the molecular pathways regulating TIMP1 and collagen-IV. Our previous publication has shown that 7-day co-culture with pericytes significantly increased the expression of insulin receptor in iBMECs. Therefore, we hypothesized that endothelial insulin signaling pathway might be activated in the co-culture model. As shown in **Figure 3B**, two major downstream targets in the insulin signaling pathway, AKT and ERK, were significantly activated in the iBMECs co-cultured with pericytes for 7 days. The mean fold change was 1.8 for pAKT and 2.4 for pERK in co-cultured versus solo-cultured iBMECs (**Figure 3E-F**).

**Figure 3.**
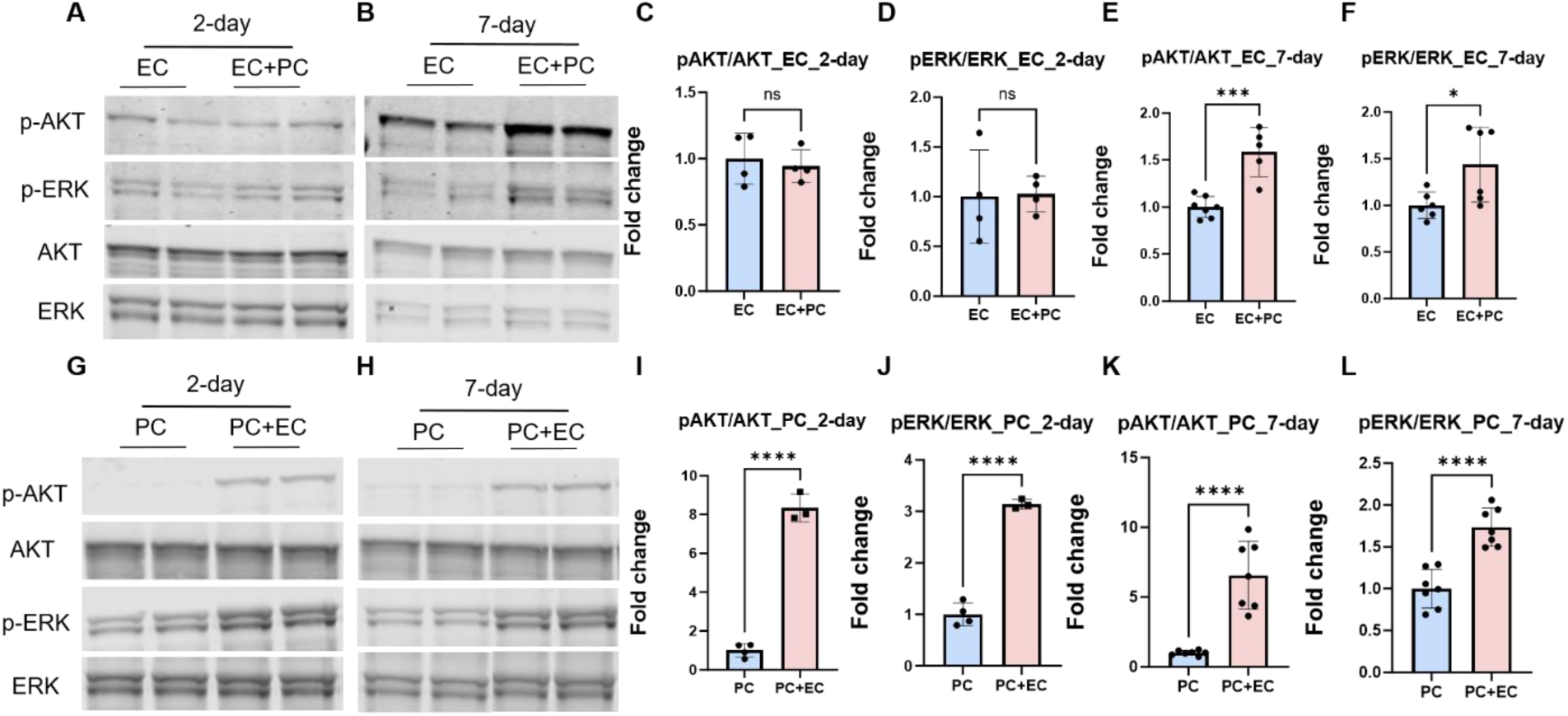
Increase in AKT and ERK phosphorylation in endothelial and pericyte co-cultures. **(A-B)** Representative immunoblots showing p-AKT, p-ERK, AKT and ERK expression in iBMECs with or without co-culture with pericytes for **(A)** 2 days and **(B)** 7 days. **(C-F)** Densitometric analysis of p-AKT and p-ERK immunoblots normalized to AKT and ERK, respectively in (A) and (B). **(G & H)** Representative immunoblots showing p-AKT, AKT, p-ERK and ERK expression in pericytes with or without co-culture with iBMECs for **(G)** 2 days and **(H)** 7 days. **(I-L)** Densitometric analysis of p-AKT and p-ERK immunoblots normalized to AKT and ERK, respectively, in (G) and (H). Data in all the bar charts were presented as fold change compared to the solo cultures. (Mean ± SD). *p<0.05, ***p<0.001, ****p<0.0001, Student’s t-test.

However, there was no significant difference in either pAKT or pERK levels between iBMECs solo cultures and those co-cultured with pericytes for 2 days (**Figure A, C-D**). In the pericytes, both 2-day and 7-day co-cultures with iBMECs showed a significant increase in AKT and ERK phosphorylation (**Figure 3G-H**). Interestingly, co-culturing for 2 days appeared to have a greater effect than 7 days in increasing pAKT and pERK expression. The pAKT expression was found to be upregulated by 8-fold in 2-day co-cultures and 6-fold in 7-day co-cultures (**Figure 3I, K**). Regarding pERK, 2-day co-cultures showed a 3-fold increase, whereas 7-day co-cultures showed a 1.7-fold increase compared to solo-cultures (**Figure 3J, L**).

### 3.4. ERK pathway regulated TIMP1 expression in BBB endothelial cells

The contribution of AKT and ERK pathways to TIMP1 expression was investigated in both hCMEC/D3 and iBMEC cultures. In hCMEC/D3 cells, it was found that insulin significantly increased TIMP1 protein expression after 15 and 30 minutes of treatment (**Figure 4A-B**). To further verify the roles of AKT versus ERK pathways, the cells were pre-treated with specific inhibitors targeting AKT (MK2206) or ERK (Trametinib), followed by insulin stimulation. As depicted in **Figure 4C-D**, inhibition of ERK pathway by trametinib significantly decreased TIMP1 expression, regardless of the presence or absence of insulin. However, AKT inhibitor MK2206 failed to show any noticeable effect. The impact of ERK inhibition on MMP activity in the endothelial cell culture media was also evaluated. Trametinib was found to elevate MMP activity in the iBMEC culture media sampled from both luminal and abluminal sides, although only the difference on the luminal side was statistically significant (**Figure 4E**). Increased MMP activity following ERK inhibition was further confirmed in the culture media of polarized hCMEC/D3 cell monolayers grown in 6-well plates. (**Figure 4F**). These results suggested that activation of the ERK pathway upregulated TIMP1 expression and inhibited downstream MMP activity in the BBB endothelial cells co-cultured with pericytes.

**Figure 4.**
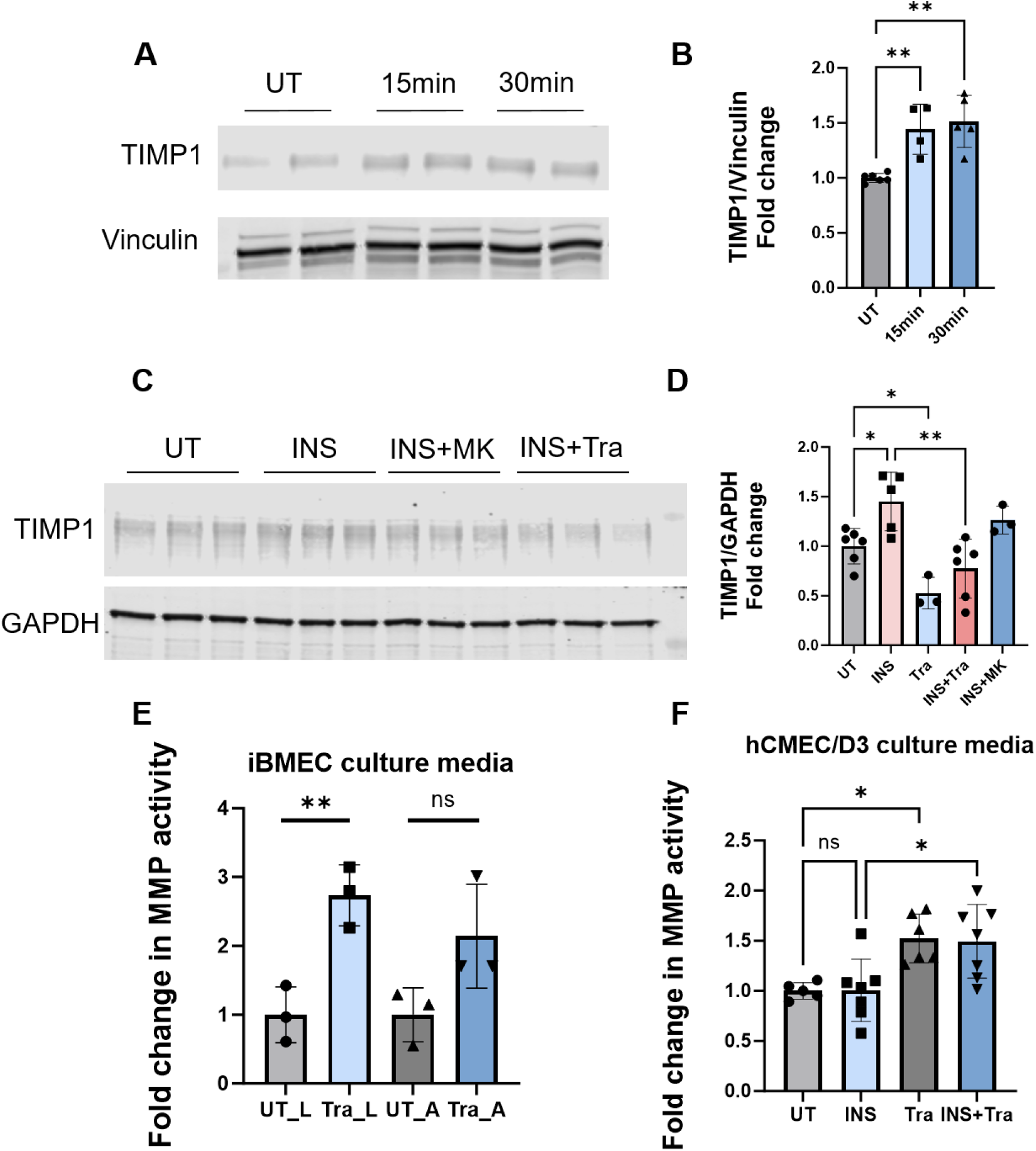
ERK pathway regulates TIMP1 expression and MMP activity in BBB endothelial cells. **(A)** Representative immunoblots depicting TIMP1 and Vinculin expression in hCMEC/D3 endothelial cells treated with 10 nM insulin for 15 and 30 minutes. **(B)** Densitometric analysis of TIMP1 immunoblots normalized to Vinculin as shown in (A). **p<0.01, one-way ANOVA with Bonferroni post-tests. **(C)** Representative immunoblots illustrating TIMP1 and GAPDH expression in hCMEC/D3 endothelial cells stimulated with 10 nM insulin with or without pretreatment of AKT inhibitor (MK2206, 10 µM) or ERK inhibitor (trametinib, 10 µM). Cells were pretreated with inhibitors for 24 hours followed by insulin stimulation for 30 minutes. **(D)** Densitometric analysis of TIMP1 immunoblots normalized to GAPDH shown in (C). *p<0.05, **p<0.01, one-way ANOVA with Bonferroni post-tests. **(E)** MMP activity in conditioned media sampled from luminal (L) and abluminal (A) sides of iBMEC endothelial monolayer treated with 10 µM trametinib for 6 hours. **p<0.01, Student’s t-test. **(F)** MMP activity in conditioned media sampled from hCMEC/D3 endothelial cell culture plates treated with insulin (10 nM) with or without pretreatment of trametinib (10 µM). Cells were pretreated with inhibitors for 24 hours followed by insulin stimulation for 30 minutes. *p<0.05, one-way ANOVA with Bonferroni post-tests. Data in all the bar charts were presented as fold change compared to untreated controls. (Mean ± SD).

### 3.5. BDNF-containing exosomes mediate the interaction between endothelial cells and pericytes

To investigate the underlying molecular mechanisms by which endothelial-pericyte interactions activate the ERK-TIMP1 signaling pathway, we treated hCMEC/D3 endothelial cells with culture media conditioned by solo- or co-cultured iBMECs.

Interestingly, the conditioned media from the luminal side of 7-day co-cultured iBMECs significantly increased TIMP1 expression in hCMEC/D3 endothelial cells compared to those treated with conditioned media from the luminal side of solo-cultured iBMECs (**Figure 5A-B**). Further, we observed an increase in ERK phosphorylation in hCMEC/D3 cells treated with conditioned media obtained from the iBMECs co-cultures compared to the solo-culture (**Figure 5A**). The difference in ERK phosphorylation following treatment of abluminal media was statistically significant (**Figure 5C**). Based on the recent reports that projected exosomes as mediators of cell-cell interactions, we hypothesized that co-culture with pericytes may enhance the endothelial secretion of biomolecules into exosomes which subsequently triggers the activation of ERK-TIMP1 pathway. To test this hypothesis, we first successfully isolated exosomes from iBMEC conditioned media. The exosomes were characterized by their particle size distribution determined by NTA (**Figure 5D**) and expression of the canonical marker CD9 as determined by western blot analysis (**Figure 5E**). Transcriptomic analysis from our previous publication has demonstrated a significant increase in BDNF gene expression in BBB endothelial cells when co-cultured with pericytes. Further, BDNF is a growth factor known to activate the ERK pathway. Therefore, we determined the BDNF protein concentration in isolated exosomes using an ELISA assay. Remarkably, the BDNF concentration in exosomes derived from the co-culture media on the abluminal side was significantly higher than the combined BDNF concentrations from iBMEC and pericyte solo-culture media (**Figure 5F)**. These results indicated that endothelial-pericyte interactions enhance BDNF secretion in exosomes into the abluminal side, potentially activating the ERK-TIMP1 signaling pathway in both cell types.

**Figure 5.**
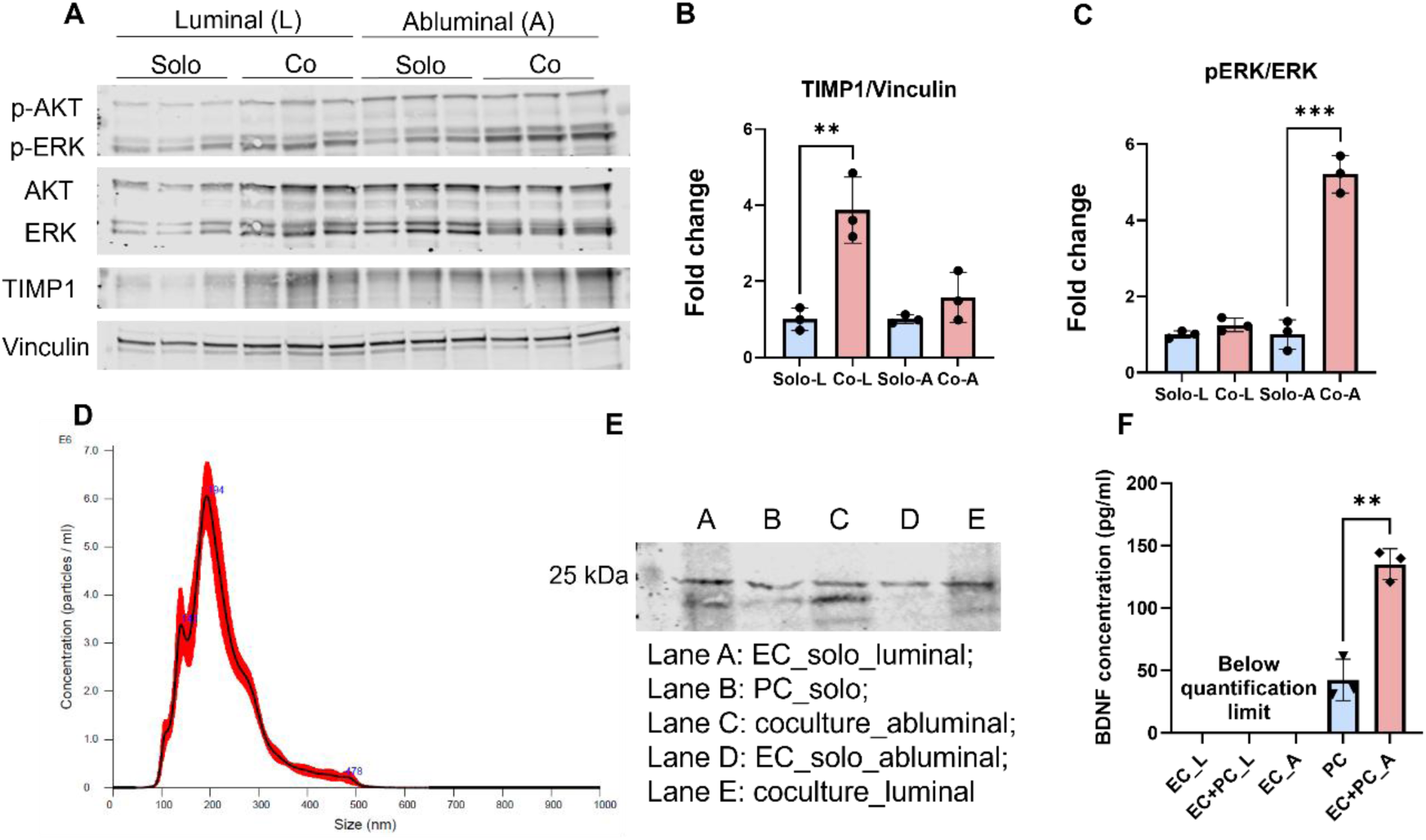
BDNF-containing exosomes mediated the interaction between endothelial cells and pericytes. **(A)** Representative immunoblots showing p-AKT, p-ERK, AKT, ERK, TIMP1 and Vinculin expression in hCMEC/D3 endothelial cells treated with conditioned media obtained from iBMECs solo- or co-cultured with pericytes on both luminal and abluminal sides. **(B-C)** Densitometric analysis of immunoblots for **(B)** TIMP1 normalized to Vinculin and **(C)** pERK normalized to ERK in Figure A. **p<0.01, ***p<0.001, one-way ANOVA with Bonferroni post-tests. **(D)** Size distribution of exosomes isolated from iBMEC conditioned media. The size was determined by NTA (red bars indicate SD, *n* = 3). **(E)** Representative immunoblots showing expression of the canonical exosome marker CD9 in exosome lysates. **(F)** BDNF concentration within exosomes derived from media of iBMEC monolayers with or without co-culture with pericytes. **p<0.01, student’s t-test.

### 3.6. Co-culture with endothelial cells decreased LRP1-mediated F-Aꞵ_42_ accumulation in pericytes

Our recent study demonstrated that inhibition of insulin signaling leads to increased Aβ42 accumulation in BBB endothelial cells[22]. Given that insulin signaling is activated in pericytes during co-culture with endothelial cells, we next examined how co-culture conditions influence Aꞵ_42_ accumulation in pericytes. As shown in Figure **6A-B**, 7-day co-culture significantly decreased F-Aβ_42_ uptake by pericytes. To understand the underlying molecular mechanisms, we examined the protein expression of putative Aβ_42_ receptors/transporters in pericytes. Our results revealed that co-culture with iBMECs for 7 days significantly reduced the expression of LRP1 (**Figure 6C**). Interestingly, RAGE expression was significantly upregulated in co-cultured pericytes. Additionally, caveolin-1 protein expression was found to be significantly reduced in pericytes co-cultured with iBMECs for 7 days. Given that caveolin-1 is crucial for caveolae formation on the plasma membrane, we explored the potential involvement of caveolae-mediated endocytosis in F-Aβ_42_ uptake by pericytes. However, no significant difference was detected in F-Aβ_42_ accumulation in pericytes treated with the caveolae-mediated endocytosis inhibitors Nystatin or Filipin (**Figure 6H-I**). These results suggested that caveolae may not contribute to F-Aβ_42_ endocytosis in pericytes and the role of caveolin-1 in modulating Aβ_42_ uptake process needs further investigation. Further, we explored the role of caveolin-1 in activating pericyte insulin signaling pathways. As shown in **Figure 6I-K**, when caveolin-1 protein was knocked down by siRNA, both AKT and ERK phosphorylation were significantly increased in pericytes.

**Figure 6.**
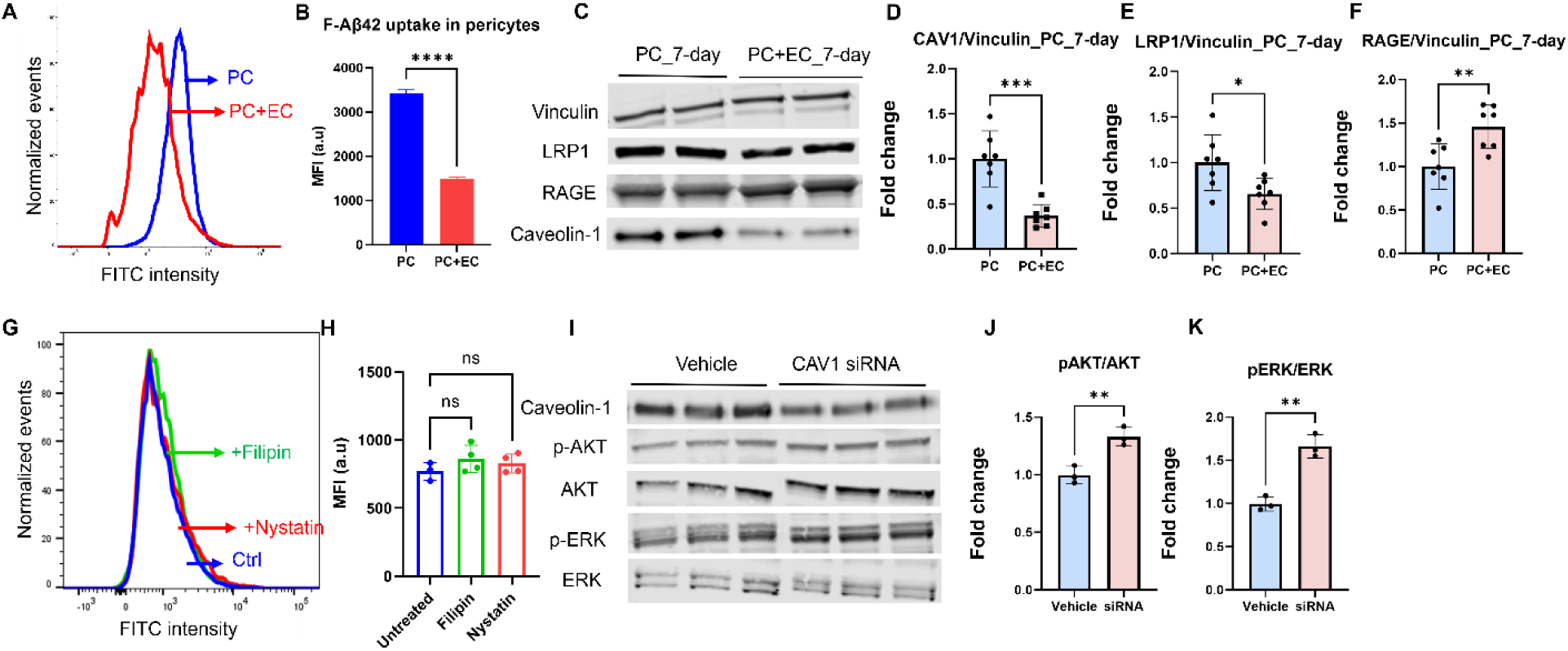
Co-culture with endothelial cells decreased LRP1-mediated endocytosis of F-Aβ_42_ in pericytes. **(A)** Representative histograms from flow cytometry study showing pericytes incubated with 1 µM FITC-labeled Aβ_42_ (F-Aβ_42_) for 1 hour. Pericytes were solo or co-cultured with endothelial cells for 7 days. On the day of the experiment, the media were aspirated, and cells were transferred to separate plates for F-Aβ_42_ incubation. **(B)** Median fluorescence intensity (MFI) of the cell population in solo and co-cultured pericytes. ****p<0.0001, unpaired two-tailed t-test. **(C)** Representative immunoblots showing caveolin-1, LRP1, RAGE, and Vinculin expression in pericytes with or without endothelial co-culture for 7 days. **(D-F)** Densitometric analysis of immunoblots for **(D)** caveolin-1, **(E)** LRP1, **(F)** RAGE normalized to Vinculin as shown in (C). **(G)** Representative histograms depicting pericytes incubated with 1 µM F-Aβ_42_ for 1 hour, with or without pretreatment of caveolae-mediated endocytosis inhibitors Nystatin and Filipin-III. **(H)** Median cellular fluorescence intensity (MFI) in untreated and inhibitor-treated pericytes. **(I)** Representative immunoblots showing caveolin-1, p-AKT, AKT, p-ERK and ERK expression in pericytes treated with siRNA targeting caveolin-1. **(J)** Densitometric analysis of p-AKT and p-ERK immunoblots normalized to AKT and ERK, respectively in Figure (I) **p<0.01, unpaired two-tailed t-test.

Pericytes play a vital role within the neurovascular unit by interacting with neighboring cells, including endothelial cells and astrocytes, to maintain the functionality of the BBB. Unfortunately, pericytes degenerate in AD which leads to BBB breakdown, contributing to both onset and progression of the disease[15]. Previous research has shown that pericytes loss is associated with increased BBB permeability, accompanied with downregulation of tight junction proteins[14,15]. However, the underlying molecular mechanisms remain unknown.

In this study, we investigated the impact of endothelial-pericytes interaction on the TIMP-MMP balance, a critical regulator of BBB integrity under various pathophysiological conditions. The key findings of the current study include: 1) co-culturing BBB endothelial cells and pericytes in a non-contact cell culture insert model significantly increased the TIMP1 and collagen-IV expression in both cell types compared to their respective solo-cultures, whereas soluble MMP enzyme activity in the co-culture media was inhibited; 2) the insulin signaling pathway was activated in both cell types in the co-culture and ERK was found to regulate TIMP1 expression in endothelial cells; 3) co-culture with endothelial cells protected pericytes against Aβ_42_ accumulation by decreasing the expression of LRP1 receptor; 4) activation of the ERK pathway in both cell types was potentially mediated by BDNF-containing exosomes in the co-culture system.

The role of MMP enzymes in AD pathogenesis is multifaceted. Both beneficial and detrimental effects of MMPs on AD pathologies have been reported. On one hand, MMPs mitigate Aβ deposition by promoting non-amyloidogenic processing of the amyloid precursor protein[23] and enzymatic degradation of Aβ[24]. On the other hand, MMPs contribute to BBB disruption by degrading tight junction and basement membrane proteins [6,7]. MMP activity is modulated by TIMPs, which are endogenous inhibitors, and a balanced TIMP/MMP ratio is instrumental to BBB integrity in various disease conditions that manifest BBB disruption. In the current study, we investigated the role of TIMP-MMP in regulating the BBB and evaluated the role of pericytes. Using a well-established in vitro BBB model formed by co-culturing iBMECs and hBVPs, known for its high transendothelial electrical resistance (TEER) values, we demonstrated that endothelial-pericyte interaction significantly increased TIMP1 expression in both cell types and inhibited MMP activity in the culture media (**Figure 1**). We focused on TIMP1 among four TIMP members because it specifically inhibits soluble MMPs that are implicated in AD including MMP2 and MMP9[25]. Decreased TIMP1 levels have been observed in the plasma of AD patients compared to healthy controls[26]. Additionally, an increased plasma MMP9/TIMP1 ratio has recently been proposed as a novel biomarker for AD diagnosis[27]. Although TIMP1 has been shown to have beneficial effects in repairing BBB disruption during focal cerebral ischemia[8,28] and intracerebral hemorrhage[9], its role in BBB disruption in the pathogenesis of AD remains unknown.

Our findings suggested that loss of endothelial-pericyte interaction disturbed the TIMP1/MMP balance, which provided a novel molecular mechanism underlying BBB disruption observed during pericyte degeneration in AD. Importantly, increased TIMP1 expression was observed in both 2-day and 7-day co-culture models. In 2-day co-cultures, we investigated how pericytes contribute to BBB maintenance, given that a matured barrier has already been established before exposure to pericytes. However, the 7-day co-culture model assessed the impact of pericytes on BBB formation. Our results indicated that pericytes are essential for both BBB formation and maintenance by upregulating TIMP1 expression. Interestingly, a previous study indicated that endothelial-pericyte interaction increased MMP9 secretion and BBB permeability in vitro[29]. However, that study investigated the pericyte impact by adding puromycin into the cell culture, which may have pleotropic effects on MMP9 secretion and confound the results. Moreover, it utilized primary endothelial cells and pericytes derived from porcine brain, which generally display lower TEER values and weaker BBB characteristics compared with the iBMECs used in the current study. Consistent with our findings, another study demonstrated that co-culturing iBMECs with pericytes enhanced TEER in stressed monolayers[30]. In that study, the stressed phenotype was induced by omitting Rho-associated coiled-coil containing protein kinase (ROCK) inhibitors, resulting in substantially reduced TEER values.

TIMP1 has been shown to protect tight junction proteins from degradation by MMPs in brain endothelial cells[31]. Yet, the influence of TIMP1 on ECM proteins, additional substrates for MMPs that are equally important but less studied with respect to BBB integrity, remains unclear. Therefore, we tested if the observed increase in TIMP1 expression during co-culture has any effect on collagen-IV levels in endothelial cells and pericytes. Our results have demonstrated that endothelial-pericyte interactions significantly increased collagen-IV expression in both cell types, regardless of the co-culture duration (**Figure 2**). Collagen-IV is the most abundant component of the basement membrane with six α chains (COL4A1-6). By deleting exon 41 of COL4A1, Jeanne et al, demonstrated that mutant collagen led to intracerebral hemorrhage and porencephaly[32]. Importantly, these effects were only observed when collagen mutation occurred in brain endothelial cells and pericytes, but not in astrocytes. Further, collagen-IV has been shown to inhibit Aβ fibril formation[33]. These findings indicated that collagen-IV has beneficial effects in preventing BBB disruption and Aβ pathology in AD. Our findings implied that pericyte degeneration could contribute to these pathologies potentially by disrupting collagen-IV in both endothelial cells and pericytes.

We next investigated the molecular pathways by which endothelial-pericyte interaction upregulated TIMP1 expression. Our recent publication has shown that pericytes activated pathways that protect endothelial cells against insulin resistance. Further, the endothelial insulin receptor protein expression was upregulated in the presence of pericytes[17]. Therefore, we hypothesized that the downstream insulin signaling pathway might be impaired when the endothelial-pericyte interaction was disrupted. Indeed, our results indicated that metabolic (AKT) and mitogenic (ERK) arms of the insulin signaling were activated in both endothelial cells and pericytes in the co-cultures (**Figure 3**). Brain endothelial cells have been demonstrated to secrete platelet-derived growth factor (PDGF)-BB, which then binds to PDGFR-β receptors on pericytes, initiating downstream signaling pathways that regulate survival, migration, proliferation and differentiation[34]. Further, a recent study found that PDGF-BB:PDGFR-β signaling activated the ERK pathway to promote pericyte proliferation, and the AKT pathway to trigger inflammatory responses[35]. The findings from current study confirmed the stimulatory effects of endothelial cells on AKT and ERK pathways in pericytes.

Furthermore, we revealed the regulatory dynamics, showing that a 2-day co-culture induced greater ERK phosphorylation compared to the 7-day co-culture. This result suggested that a matured endothelial barrier might exert a greater impact on pericyte proliferation. In the pericyte 7-day co-cultures, we also observed that caveolin-1 protein expression was significantly reduced. More importantly, a separate study was conducted to knock down caveolin-1 and it was shown to increase the AKT and ERK phosphorylation (**Figure 6I-K**). The inhibitory effect of caveolin-1 on ERK phosphorylation has been reported in vascular smooth muscle cells, another type of mural cells[36]. These findings suggest a potential molecular mechanism whereby endothelial-pericyte interactions may enhance insulin signaling in pericytes by downregulating caveolin-1 expression. Conversely, the influence of pericytes on endothelial ERK and AKT pathways has been less explored compared to the well-documented effects of endothelial cells on pericytes. One previous study demonstrated that retinal pericytes activated ERK pathway in brain endothelial cells in a co-culture model, which is consistent with the current findings using BBB pericytes[37].

Interestingly, this study found activation of the protein kinase C (PKC) pathway in endothelial cells co-cultured with pericytes, whereas our results demonstrated activation of the protein kinase B (AKT) pathway. This suggests a subtle difference between retinal and brain endothelial-pericyte interactions, which may maintain distinct phycological and functional regulation.

We also investigated to determine if the increased expression of TIMP1 in co-cultured endothelial cells was a downstream effect of ERK or AKT activation. Our results suggested that insulin directly increased TIMP1 expression in hCMEC/D3 cells (**Figure 4A**). Although hCMEC/D3 monolayers exhibit weaker tight junctions compared with iBMECs, they serve as an appropriate *in vitro* model for this study, as the focus is on signaling pathways rather than permeability. Similar effects were observed in adipocytes, in which insulin increased the levels of TIMP1 mRNA[38]. These results were consistent with our previous transcriptomic analysis in which insulin activated inhibition of matrix metalloproteinase pathway and increased gene expression of all the TIMP family members in BBB endothelial cells[16]. Importantly, we further showed that trametinib blocked the insulin effect on TIMP1 expression, whereas MK2206 had no impact (**Figure 4C**). These results demonstrated that ERK, instead of AKT activation was responsible for the upregulation of TIMP1 expression in endothelial cells. The ERK pathway has been reported to mediate the effects of TNF-α and TGF-β in increasing TIMP1 expression in trabecular meshwork[39] and fibrosarcoma cells[40]. In addition, previous research have shown that focal ischemia activated ERK enhanced both MMP9 and TIMP1 expression in cerebrovascular microvessels[41]. These findings suggest that the endothelial-pericyte interactions increased endothelial TIMP1 expression via activation of the ERK pathway. Although TIMP1 upregulation and ERK activation were also observed in pericytes, the causal relationship needs to be further characterized. It is important to note that activation of the ERK and AKT pathways may exert other biological effects in BBB endothelial cells and pericytes. Insulin resistance, which is characterized by disruption of ERK and AKT pathways in insulin-responsive cells, is closely associated with AD according to both epidemiological and pathophysiological evidence[42,43]. BBB insulin resistance is of particular importance because our previous publication has shown that endothelial insulin signaling pathway regulates various BBB functions including vesicular trafficking, inflammatory regulation, as well as paracellular and transcellular permeability[16]. The current study suggested that pericyte degeneration in AD may induce insulin resistance in both endothelial cells and pericytes, ultimately leading to BBB dysfunction that exacerbates AD progression.

Our above findings indicate that endothelial-pericyte interactions activate the insulin signaling pathway, regulating TIMP1 and collagen-IV expression in both cell types. However, the specific molecular mediators responsible for triggering the ERK and AKT pathways have yet to be identified. Our recently published study has shown that brain derived nerve growth factor (BDNF) gene expression was significantly increased in endothelial cells co-cultured with pericytes[17]. Moreover, a previous study suggested a significant enhancement in BDNF secretion by pericytes in response to PDGF-BB[44]. BDNF has been shown to activate both the ERK and AKT pathways in human umbilical vein endothelial cells[45]. Given these findings, we hypothesized that BDNF secretion is elevated in the co-cultures to activate the ERK and AKT pathways in endothelial cells.

Further, we speculated that the secreted BDNF was encapsulated within exosomes because exosomes have been shown to mediate cell-cell interactions by transporting signaling molecules such as growth factors[46,47]. This hypothesis was supported by the results that BDNF concentration in the co-culture media-derived exosomes was significantly higher than that in the solo-culture (**Figure 5**). Notably, the difference in BDNF concentration was observed exclusively in exosomes isolated from the abluminal media, with no significant change on the luminal side. This finding is consistent with the observed increase in ERK phosphorylation in hCMEC/D3 cells treated with media conditioned by co-cultured iBMECs on the abluminal side. These results align with the physiological role of BDNF, which primarily supports the survival and growth of neurons located on the abluminal side. Taken together, we have shown that BBB pericytes specifically interact with BBB endothelial cells in activating ERK and AKT pathways, which is mediated by BDNF-containing exosomes.

Finally, we demonstrated that F-Aβ_42_ accumulation in the pericytes was significantly reduced in presence of BBB endothelial cells. It was previously shown that pericytes internalized Aβ_42_ peptides in AD patients and animal models, contributing to Aβ clearance from the brain[48]. However, excessive Aβ build up could lead to pericytes death due to cytotoxicity of Aβ[49]. Moreover, it was found that Aβ constricted capillaries by evoking pericytes contraction, which caused hypoperfusion observed in AD brain[13]. Our results implied that the interaction between pericytes and endothelial cells might protect pericytes from Aβ_42_-induced toxicity and potentially rescue cerebral blood flow impairment. To investigate the mechanisms underlying reduced Aβ accumulation in pericytes, we evaluated the protein expression of receptors that are implicated in Aβ uptake in brain endothelial cells. Interestingly, while RAGE expression was upregulated, significant reduction in LRP1 expression was observed (**Figure 6E-F**). These results implied that LRP1 might play a dominating role in causing reduction of Aβ accumulation in pericytes when co-cultured with endothelial cells. Indeed, it has been shown that pericytes internalized Aβ_42_ via LRP1 while the contribution of RAGE remains unknown[48]. Further, we observed a significant reduction in caveolin-1 expression in pericytes co-cultured with endothelial cells. Caveolin-1 is a major constituent of caveolae in plasma membrane and our previous publication showed that Aβ_42_ endocytosis in brain endothelial cells is caveolae mediated[50]. Therefore, we hypothesized that downregulation of caveolin-1 could disrupt the caveolae-mediated endocytosis, contributing to reduced F-Aβ_42_ accumulation in pericytes. However, the flow cytometry analysis suggested that F-Aβ_42_ uptake in pericytes was caveolae independent as neither nystatin nor filipin-III had any effect on F-Aβ_42_ accumulation.

These results further supported the notion that RAGE may play a minor role in mediating Aβ_42_ uptake by pericytes because RAGE mediated endocytosis was shown to be dependent on caveolae[51] whereas LRP1-mediated endocytosis of Aβ was associated with clathrin[52].

In conclusion, this study demonstrated that the interaction between endothelial cells and pericytes enhanced BBB integrity by upregulating collagen-IV expression in both cell types via TIMP1 and protecting pericytes against Aβ_42_ accumulation via the LRP1 receptor. Further, the insulin signaling pathways were activated in co-cultures of both cell types, and the ERK pathway was identified as a key mediator of endothelial TIMP1 upregulation driven by pericyte interactions. The proposed pathway summarizing the impact of endothelia-pericyte interactions on both cell types was presented in **Figure 7**. These findings provided mechanistic insights into BBB disruption due to pericyte degeneration and underscored the crucial role of BBB insulin resistance in contributing to cerebrovascular dysfunction in AD.

**Figure 7.**
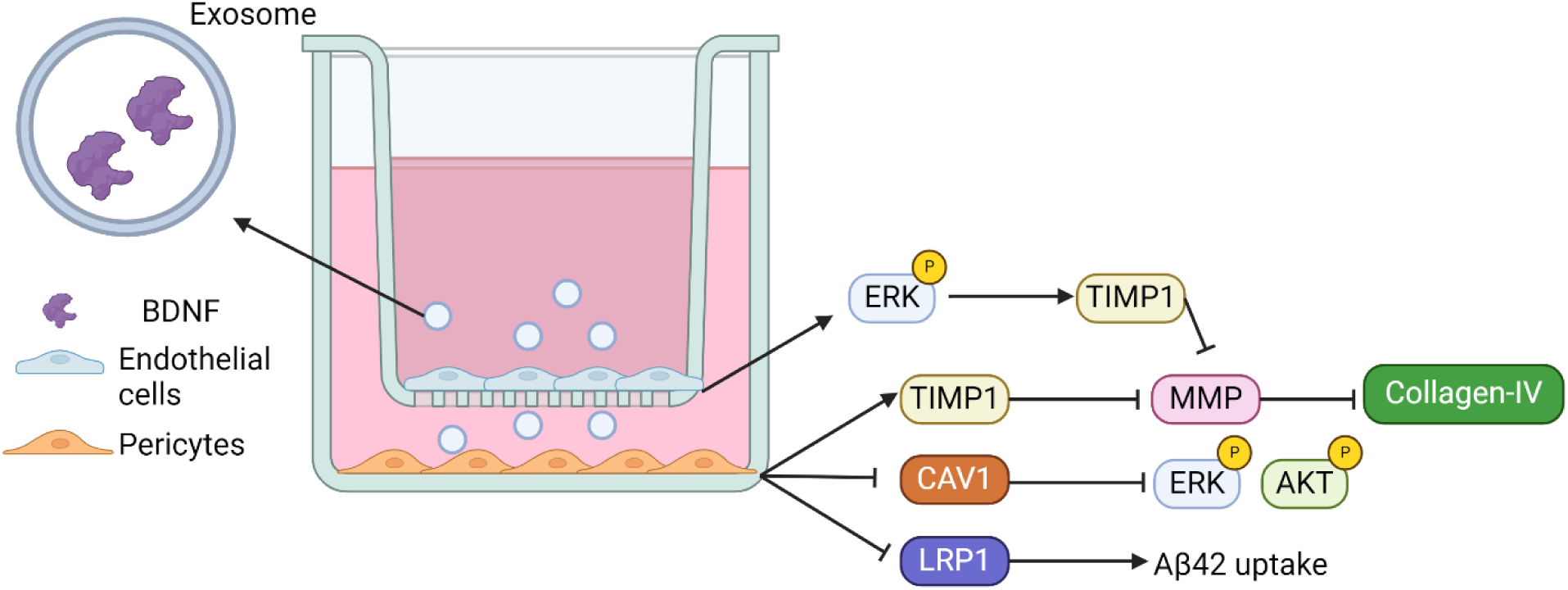
Proposed pathway summarizing the impact of endothelia-pericyte interactions on both cell types

## CRediT authorship contribution statement

**Zengtao Wang:** Writing – review & editing, Writing – original draft, Visualization, Validation, Methodology, Investigation, Formal analysis, Conceptualization. **Vaishnavi Veerareddy**: Validation, Visualization, Methodology, Investigation, Formal analysis, Writing – review & editing. **Paulina M. Eberts**: Resources, Methodology, Writing - Review & Editing. **Samira M. Azarin**: Resources, Methodology, Writing - Review & Editing. **Karunya K. Kandimalla**: Writing – review & editing, Funding acquisition, Supervision, Conceptualization.

## Data availability

All data generated or analyzed during this study are included in this published article.

